# Aging affects K_V_7 channels and perivascular-adipose tissue-mediated vascular tone

**DOI:** 10.1101/2021.04.11.438975

**Authors:** Yibin Wang, Fatima Yildiz, Andrey Struve, Mario Kassmann, Friedrich C. Luft, Maik Gollasch, Dmitry Tsvetkov

## Abstract

Aging is an independent risk factor for hypertension, cardiovascular morbidity, and mortality. However, detailed mechanisms linking aging to cardiovascular disease are unclear. We studied the aging effects on the role of perivascular adipose tissue and downstream vasoconstriction targets, voltage-dependent K_V_7 channels, and their pharmacological modulators (flupirtine, retigabine, QO58, QO58-lysine) in a murine model. We assessed vascular function of young and old mesenteric arteries *in vitro* using wire myography. We also performed bulk RNA sequencing and quantitative reverse transcription polymerase chain reaction tests in mesenteric arteries and perivascular adipose tissue to elucidate molecular underpinnings of age-related phenotypes. Results revealed impaired perivascular adipose tissue-mediated control of vascular tone particularly via K_V_7.3-5 channels with increased age through metabolic and inflammatory processes and release of perivascular adipose tissue-derived relaxation factors. Moreover, QO58 was identified as novel pharmacological vasodilator to activate XE991-sensitive KCNQ channels in old mesenteric arteries. Our data suggest that targeting inflammation and metabolism in perivascular adipose tissue could represent novel approaches to restore vascular function during aging. Furthermore, QO58 represents a novel tool for cardiovascular and hypertension research in aging.

## INTRODUCTION

Hypertension is the leading risk factor of death worldwide, especially for persons aged 50–74 years and 75 years and older (Collaborators, 2020). Human life expectancy promises to increase steadily, which will further amplify age-related effects (Vollset et al., 2020). Rigorous research is necessary to address these challenges. Despite remarkable progress in understanding molecular biology of aging, detailed mechanisms linking aging to cardiovascular disease are still unclear (North & Sinclair, 2012). Mice are a utilitarian model to investigate age-related effects (DuPont et al., 2016). Our past studies have shed light on the regulation of vasculature tone by perivascular adipose tissue (PVAT) (Gollasch, 2017). Fatty tissue surrounding blood vessels is now recognized as an integral endocrine/paracrine organ. In addition to the endothelium, PVAT releases vasoactive compounds to cause relaxation of blood vessels known as the anti-contractile effect of PVAT (Lohn et al., 2002). PVAT relaxation factors (PVATRFs) have been proposed and such factors could be pivotal in aging. Our earlier work suggests that PVAT paracrine effects are caused by opening of potassium (K^+^) channels in vascular smooth muscle cells (Verlohren et al., 2004). The KCNQ-type, K_V_7 channels represent the most likely candidates as largely supported by studies with XE991, a highly effective blocker of these channels (Schleifenbaum et al., 2010) (Tsvetkov et al., 2017). In fact, the K_V_7 family represents a new target for hypertension treatment (Schleifenbaum et al., 2010) (Jepps et al., 2011) (Mani et al., 2016). K_V_7 channel function determines sensitivity to key regulators of coronary tone in diabetes, which expands therapeutic potential even further (Morales-Cano et al., 2015) (Barrese, Stott, & Greenwood, 2018). However, toxicity issues of currently available K_V_7 channel modulators, such as retigabine and flupirtine, has hampered drug development directed at this target (European.Medicines.Agency, 2018) (FDA, 2013). Pyrazolo[1,5-a]pyrimidin-7(4H)-one compounds (e.g. QO58) have been identified as novel KCNQ channel openers, which can cause remarkable leftward shifts of voltage-dependent activation of K_V_7 channels (Jia et al., 2011). Newly emerging RNA sequencing technologies coupled with established techniques could enable researching these new compounds (Tabula Muris, 2020). We hypothesize that age could affect the function of PVAT as mediated by K_V_7 channels. We employed the established flupirtine and retigabine, as well as novel compounds (QO58 and QO58-lysine) as K_V_7 channel activators in isolated mesenteric arteries from young and old mice.

## RESULTS

### Aging impairs PVAT-mediated control of vascular tone

First, we examined the role of aging in the anti-contractile effects of PVAT. Isolated mesenteric arteries were contracted by alpha1 adrenoceptor (alpha1-AR) stimulation with methoxamine (ME). To test whether or not PVAT regulation on the arterial tone is impaired in aging we performed a series of experiments using arteries from young (3 months old), 12-, and 16-months old mice (Figure 1). Arteries were prepared either with (+) PVAT or without (−) PVAT. Mesenteric artery rings of young mice displayed strong anti-contractile effects of PVAT, namely the dose-response curve for vasocontractions of (+) PVAT rings by ME was shifted to the right, compared to (–) PVAT rings (Figures 1 A, B). In contrast, (–) PVAT artery rings from 1 year old mice displayed contractions in response to alpha1-AR agonist similar to (+) PVAT rings (Figures 1 C, D). To substantiate the results, we performed similar experiments using artery rings isolated from 16-months old mice. Alpha1-AR agonist induced contractions were similar between (−) PVAT rings and (+) PVAT rings (Figures 1 E, F). Together, the results suggest that the anti-contractile effects of PVAT are impaired in aging.

**Figure 1.**
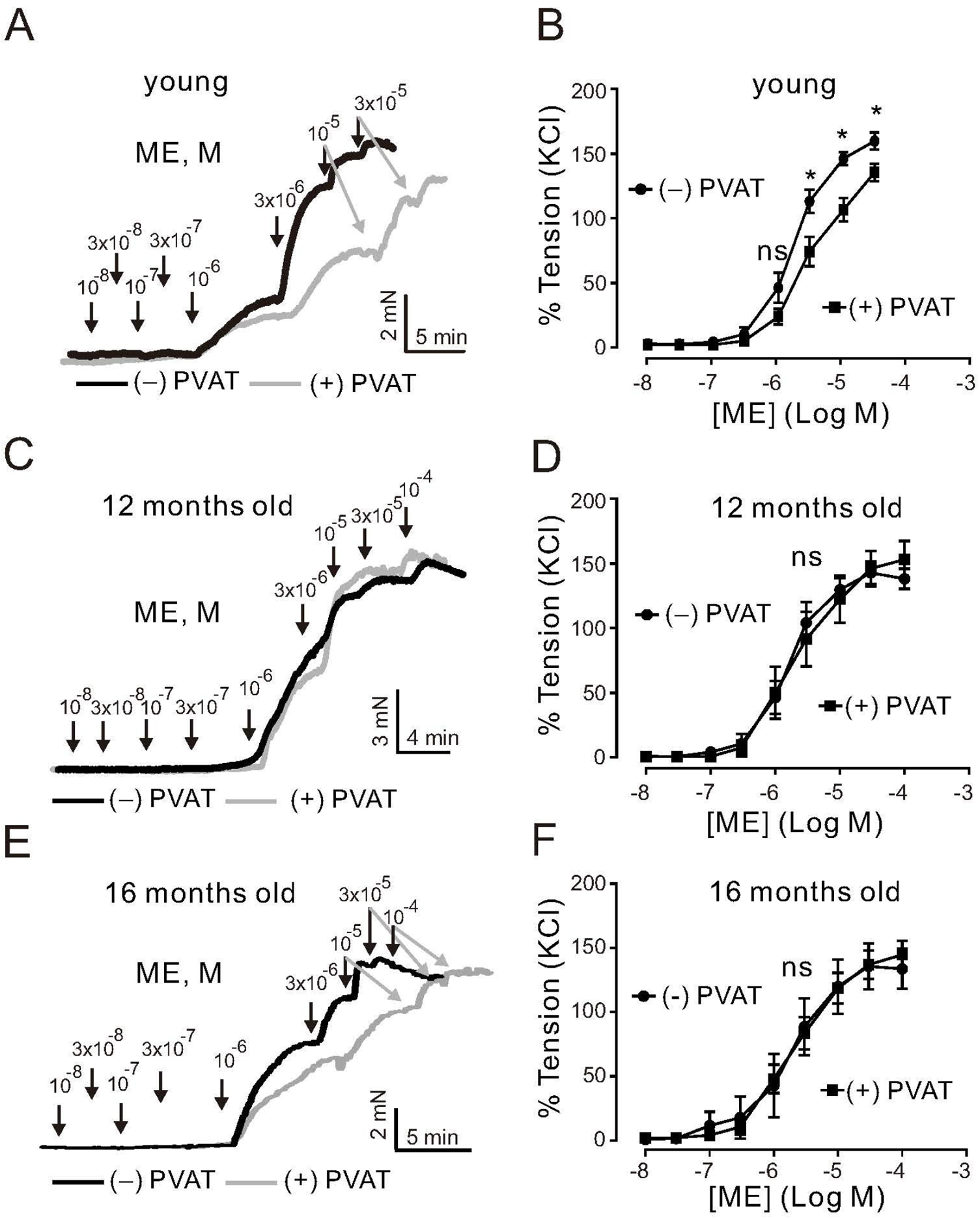
Effects of aging on regulation of arterial tone by a1-agonists Methoxamine (ME) and perivascular adipose tissue (PVAT). (A) Original traces showing a1-agonist-induced contractions in (−) PVAT and (+) PVAT mesenteric artery rings isolated from young mice. (B) Concentration-response relationships for a1-agonist-induced contractions in (−) PVAT (n = 10) or (+) PVAT (n = 10) mesenteric arteries from young animals. (C) Original traces showing aging effects on a1-agonist-induced contractions in (−) PVAT and (+) PVAT mesenteric artery rings isolated from 12-months old mice. (D) Concentration-response relationships for a1-agonist-induced contractions of (+) PVAT (n = 6) and (−) PVAT (n = 9) artery rings isolated from 12-months old mice. (E) Original traces showing aging effects on a1-agonist-induced contractions in (−) PVAT and (+) PVAT mesenteric artery rings isolated from 16-months old mice. (F) Cumulative concentration-response relationships to a1-agonist in (−) PVAT (n = 7) and (+) PVAT (n = 9) mesenteric arteries in 16-months old mice. *p < 0.05. Two-way ANOVA followed by Bonferroni post hoc test. Data are mean and SEM.

### K_V_7 channel function in PVAT is affected by age

Next, we assessed the role of K_V_7 channels during aging. K_V_7 channels were activated by flupirtine and retigabine, which are considered as potent KCNQ3-5 activators in vascular smooth muscle (Tsvetkov et al., 2017). Flupirtine produced dose-dependent relaxations; however, the effects were reduced by increased age. For instance, in arterial rings from 12- and 24-months old mice the effects were clearly age-dependent (Figures 2 A,B). The 95% CI for EC_50_ of young, 12-, and 24-months old mice rings were 0.6 - 0.8 μM, 1.8 – 4.8 μM, and 12.4 – 43.3 μM, respectively. Retigabine caused similar effects (Figures 2 C, D).

**Figure 2.**
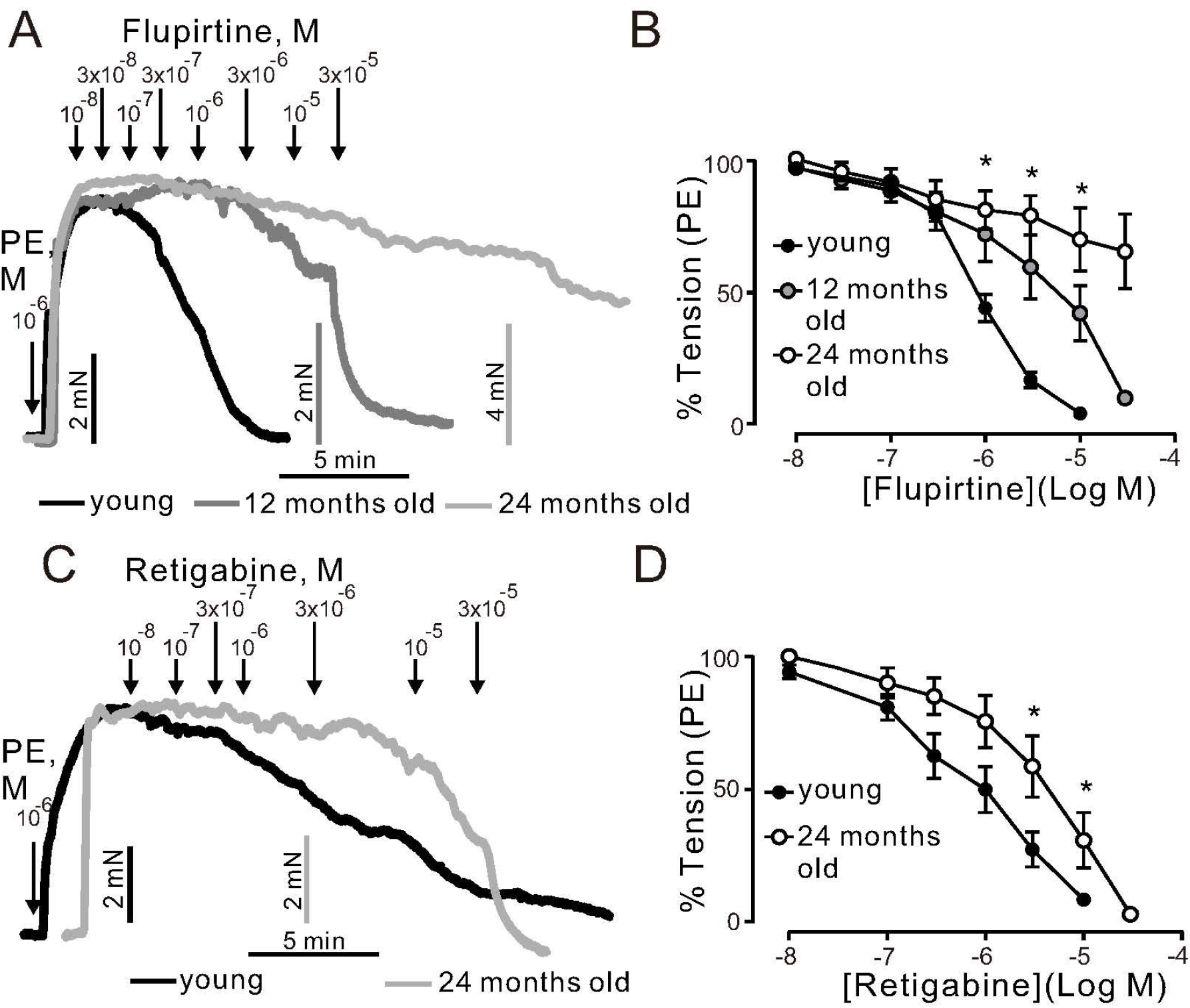
Relaxation of (−) PVAT mesenteric artery rings by KCNQ channels activators flupirtine and retigabine. (A) Representative traces showing the relaxation induced by 0.01-30 μM flupirtine in rings isolated from young, 12-months, and 24-months old mice. Mesenteric arteries were precontracted by 1 μM phenylephrine (PE). (B) Concentration-response relationships for flupirtine-induced relaxation in mesenteric arteries in young (n = 12), 12-months old (n = 10), and 24-months old (n = 9) mice. (C) Representative traces showing the relaxation induced by 0.01-30 μM retigabine in rings isolated from young, 12-, and 24-months old mice. (D) Concentration-response relationships for retigabine-induced relaxation in mesenteric arteries in young (n = 12), 12-(n = 9), and 24-months old (n = 7) mice. *p < 0.05. Two-way ANOVA followed by Tukey post hoc test. Data are mean and SEM.

We tested two novel K_V_7 channel activators, namely QO58 and QO58-lysine. QO58 produced dose-dependent relaxations. The effects were abolished by 3 µM XE991 (pan K_V_7 channel blocker) at low QO58 concentrations (<1 µM) (Figures 3 A, B). In contrast, XE991 was unable to inhibit relaxations induced by QO58-lysine (Figures 3 C, D). The data suggest that OQ58 but not QO58-lysine is capable of producing arterial relaxations through activation of XE991 sensitive KCNQ channels. Similar to flupirtine and retigabine, aging attenuated QO58-induced relaxations (Figure 3E). Thus, our data indicate that KCNQ channel function is impaired in aging.

Then, we determined whether or not the effects of flupirtine, retigabine, Q58,QO58-lysine rely on K^+^ channel activation. Raising external [K^+^] to 60 mM would be expected to diminish the effects of any K^+^ channel opener by substantially reducing the difference between the potassium equilibrium potential and membrane potential. In these conditions, contractions are primarily caused by Ca^2+^ influx through L type Ca_V_1.2 channels resistant to K^+^ channel openers (Essin et al., 2007). We found that flupirtine, retigabine, QO58, QO58-lysine produced moderate relaxation only at relatively high (≥30 µM) concentrations (Figures 4 A, B, C, D). Vehicle application produced no relaxations (Figures 4 E, F). Therefore, all four KCNQ channel activators may have off target effects on Ca^2+^ channels, namely L-type Ca_V_1.2 channels, only at higher concentrations (≥ 30 µM).

**Figure 3.**
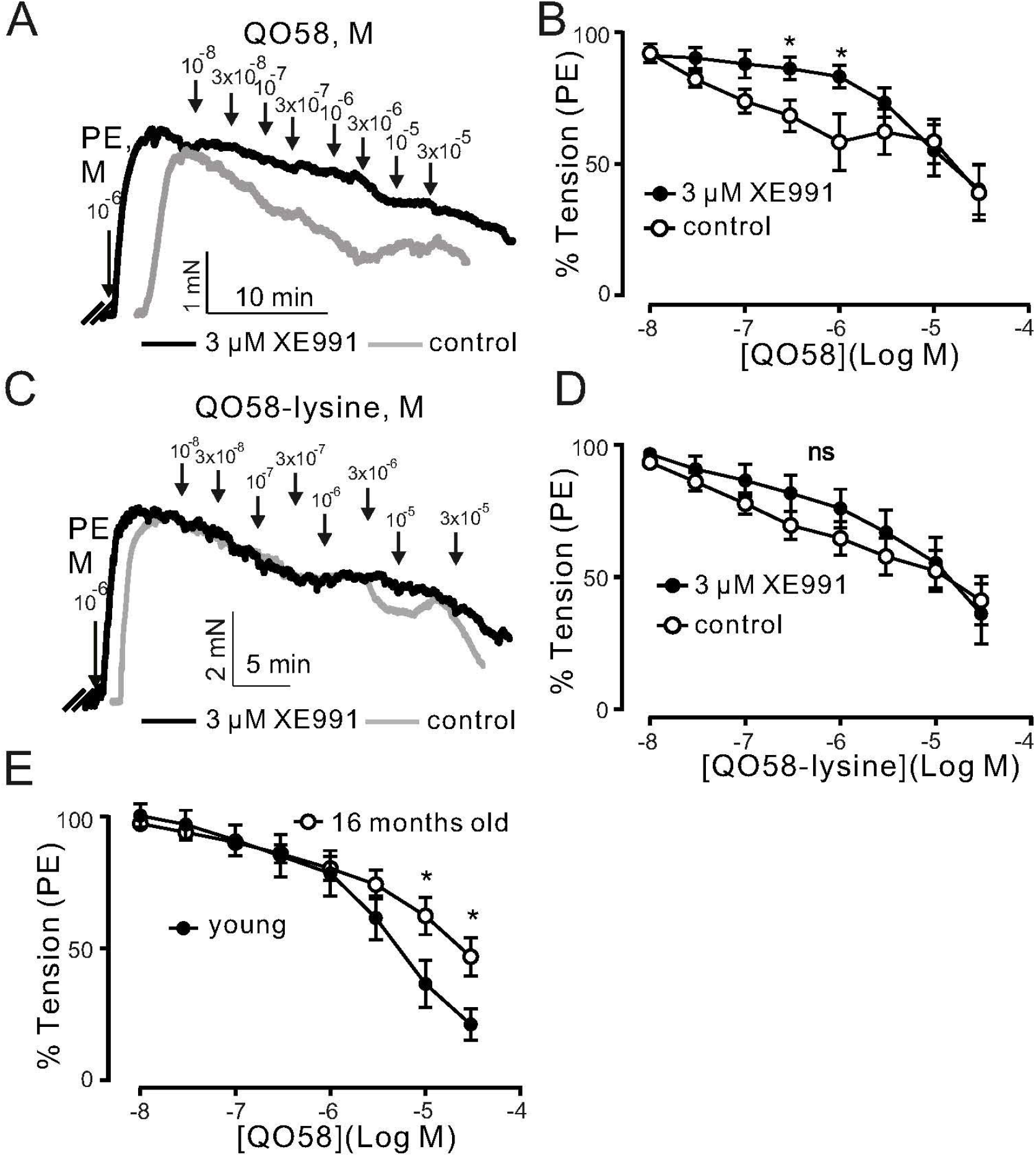
Relaxation of (−) PVAT mesenteric artery rings by novel KCNQ channel openers QO58 and QO58-lysine. (A) Original traces showing the effects of 3 μM XE991 on QO58-induced relaxation in (−) PVAT mesenteric artery rings compared with control rings without XE991. (B) Concentration-response relationships for QO58-induced relaxation in (−) PVAT mesenteric arteries from young wild-type animals after pre-incubation with 3 μM XE991 or in the absence of XE991. (C) Original traces showing the effects of 3 μM XE991 on QO58-lysine-induced relaxation in (−) PVAT mesenteric artery rings compared with control rings without XE991. (D) Concentration-response relationships for QO58-lysine-induced relaxation in (−) PVAT mesenteric arteries from young animals after pre-incubation with 3 μM XE991 or in the absence of XE991. (E) Concentration-response relationships for QO58-induced relaxation in mesenteric arteries in young (n = 11) and 16-months old (n = 13) mice. *p < 0.05. One sample t-test. Data are mean and SEM.

**Figure 4.**
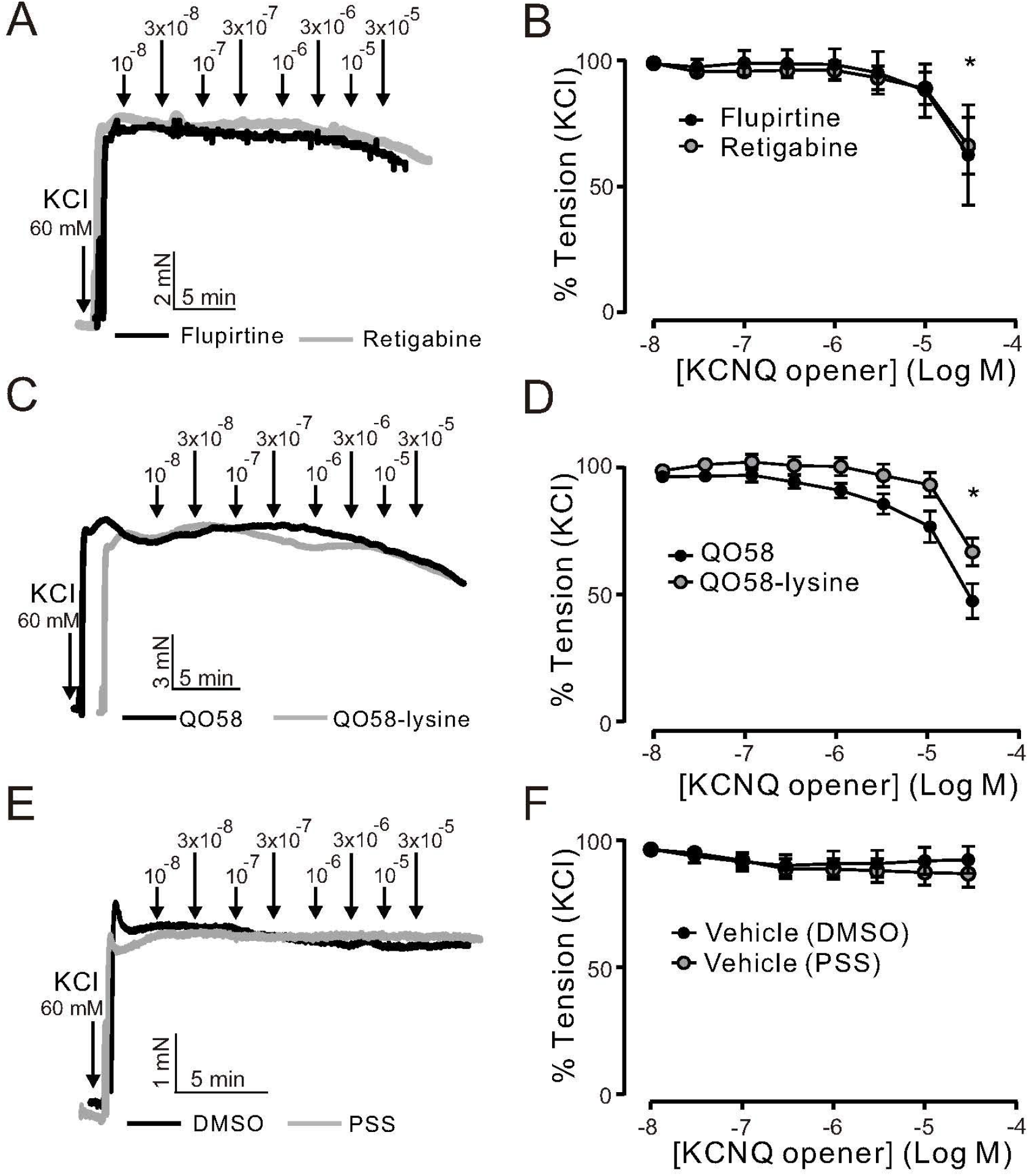
KCNQ channel openers effects on KCl-induced contraction in (−) PVAT mesenteric artery rings. (A) Original recordings showing the effects of 0.01-30 μM retigabine, 0.01-30 μM flupirtine on arterial tone of isolated mesenteric artery rings without (–) PVAT. Vessels were precontracted with 60 mM KCl. (B) Concentration-response relationships for flupirtine-(n = 6) and retigabine-induced relaxation (n = 7) in (−) PVAT mesenteric arteries from young wild-type animals. (C) Original recordings showing the effects of 0.01-30 μM QO58 and 0.01-30 μM QO58-lysine on arterial tone of isolated mesenteric artery rings without (–) PVAT. (D) Concentration-response relationships for QO58 (n = 9) and QO58-lysine-induced relaxation (n = 8) in (−) PVAT mesenteric arteries from young wild-type animals. (E) Original recordings showing the effects of vehicle (DMSO or PSS) on arterial tone of isolated mesenteric artery rings without (–) PVAT. (F) Concentration-response relationships for DMSO (n = 5) and PSS (n = 5) in (−) PVAT mesenteric arteries from young wild-type animals. *p < 0.05. paired sample t-test. Data are mean and SEM.

### RNA-Sequencing

To examine age-related changes in mRNA expression in mesenteric arteries and PVAT we performed targeted and bulk RNA sequencing (RNA-seq) utilizing arterial tissue from young and old mice. Per sample, we obtained 23 ± 2.5 million reads. ∼97,5 % of all reads were mapped to the reference mouse genome (ensembl_mus_musculus_grcm38_p6_gca_000001635_8). The principal component analysis (PCA) demonstrated tight clustering within each group and transcriptome difference between groups (Figure 5 A). In (–) PVAT mesenteric arteries isolated from 12- 16-, and 24-months old mice, we were interested in candidate genes involved in pathways regulating KCNQ channels. Figure 5 B shows the results. The data show that none of them was affected by aging. However, we found that transcripts of several ion channels were up or down-regulated in (–) PVAT mesenteric arteries during aging. The results are shown in Figure 5 C, D, E. Of note, the mRNA expression of *Kcnq1,3,4,5* was normal across the different ages. We also confirmed these results using qPCR (Figures S1 A, B, C, D). In PVAT from 12-months old mice, 2202 transcripts were upregulated and 1767 were downregulated (Figure 5 A). Top 5 down- and upregulated genes are depicted on Figure 5 G.

**Figure 5.**
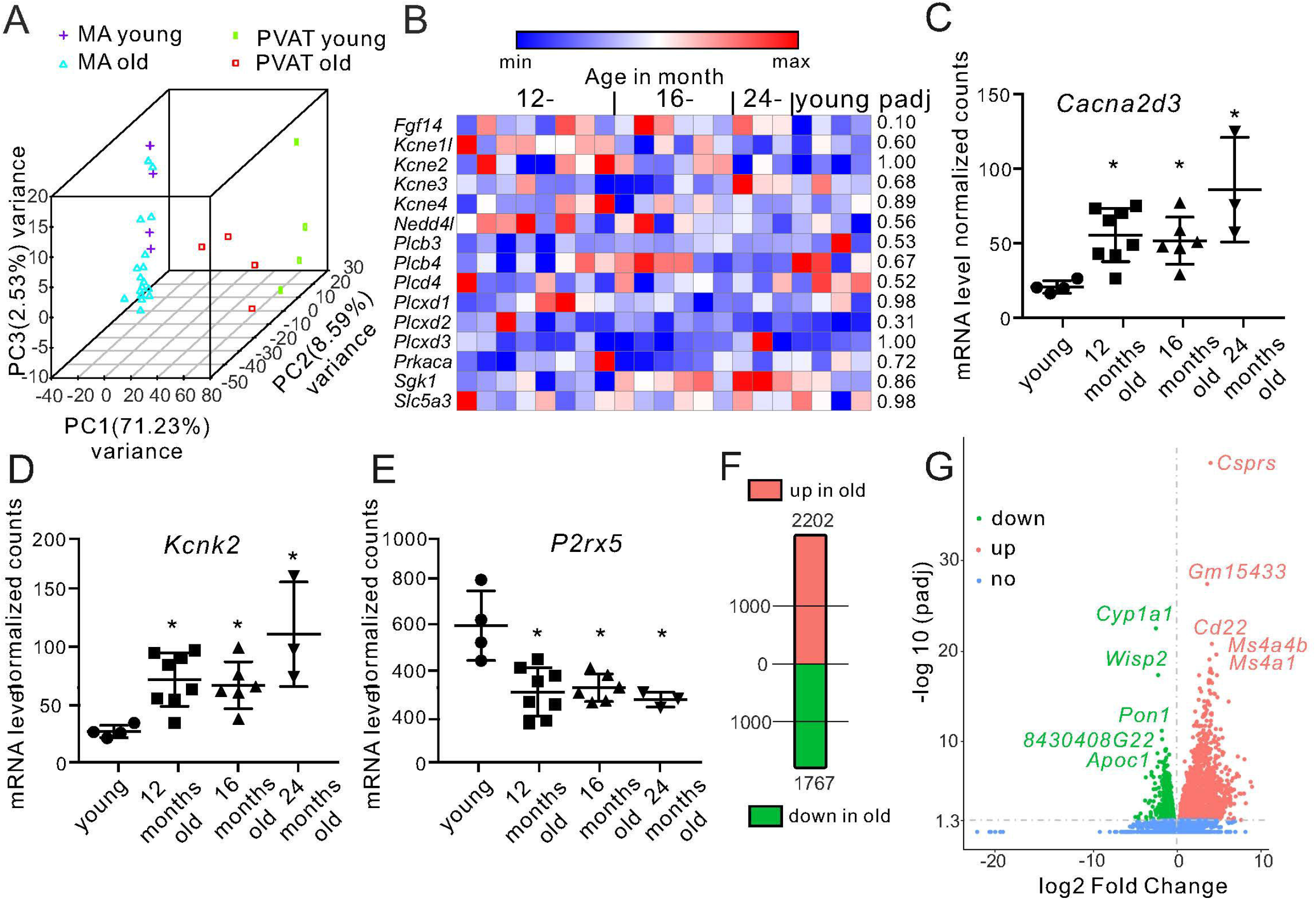
Differential gene expression in mesenteric arteries and PVAT from young and old mice. (A) Principal component analysis in samples isolated from young vs old animals. (B) Heatmap showing the expression of candidate genes involved in pathways regulating KCNQ channels (n = 8 for 12-months old; n = 6 for 16-months old; n = 3 for 24-months old; n = 4 for young mice). Data are fragments per kilobase per million mapped fragments (FPKM). (C) Relative mRNA levels for *Cacna2d3* (n = 4 for young; n = 8 for 12-months old; n = 6 for 16-months old; n = 3 for 24-months old mice) (D) Relative mRNA levels for *Kcnk2* (n = 4 for young; n = 8 for 12-months old; n = 6 for 16-months old; n = 3 for 24-months old mice) (E) Relative mRNA levels for *P2rx5* (n = 4 for young; n = 8 for 12-months old; n = 6 for 16-months old; n = 3 for 24-months old mice). (F) Number of differentially expressed genes in PVAT isolated from mesenteric arteries in young and old mice. (G) Volcano plot displaying statistical significance (adjusted p value) versus magnitude of change (fold change) for differently expressed genes in PVAT isolated from 12-months old (n=4) vs young mice (n=4). Top 5 differentially expressed genes are marked. *P < 0.05. Wald test. Data are mean and SD. Abbreviations can be found in Table S3.

### Metabolic and inflammatory pathways

Next, we performed Gene Ontology (GO) enrichment analysis using biological process (BP) terms and Kyoto Encyclopedia of Genes and Genomes (KEGG) pathways. Our data show that aged PVAT exhibited upregulated pathways associated with inflammatory processes (e.g., GO:0002250, GO:0051249, GO:0002764; mmu05150, mmu05152, mmu04060) (Table S1). Downregulated were mostly BP and pathways related to generation of precursor metabolites and energy (e.g., GO:0006091, GO:0051186, GO:0006119, mmu00190, mmu01212, mmu03320) (Table S2). In detail, the downregulated genes include mitochondrial genes associated to Parkinson (mmu05012) and Huntington (mmu05016), fatty acid metabolism (mmu01212), biosynthesis of unsaturated fatty acids (mmu01040), fatty acid elongation (mmu00062), insulin signaling (mmu04910), and PPAR pathway (mmu03320) (Figure 6).

**Figure 6.**
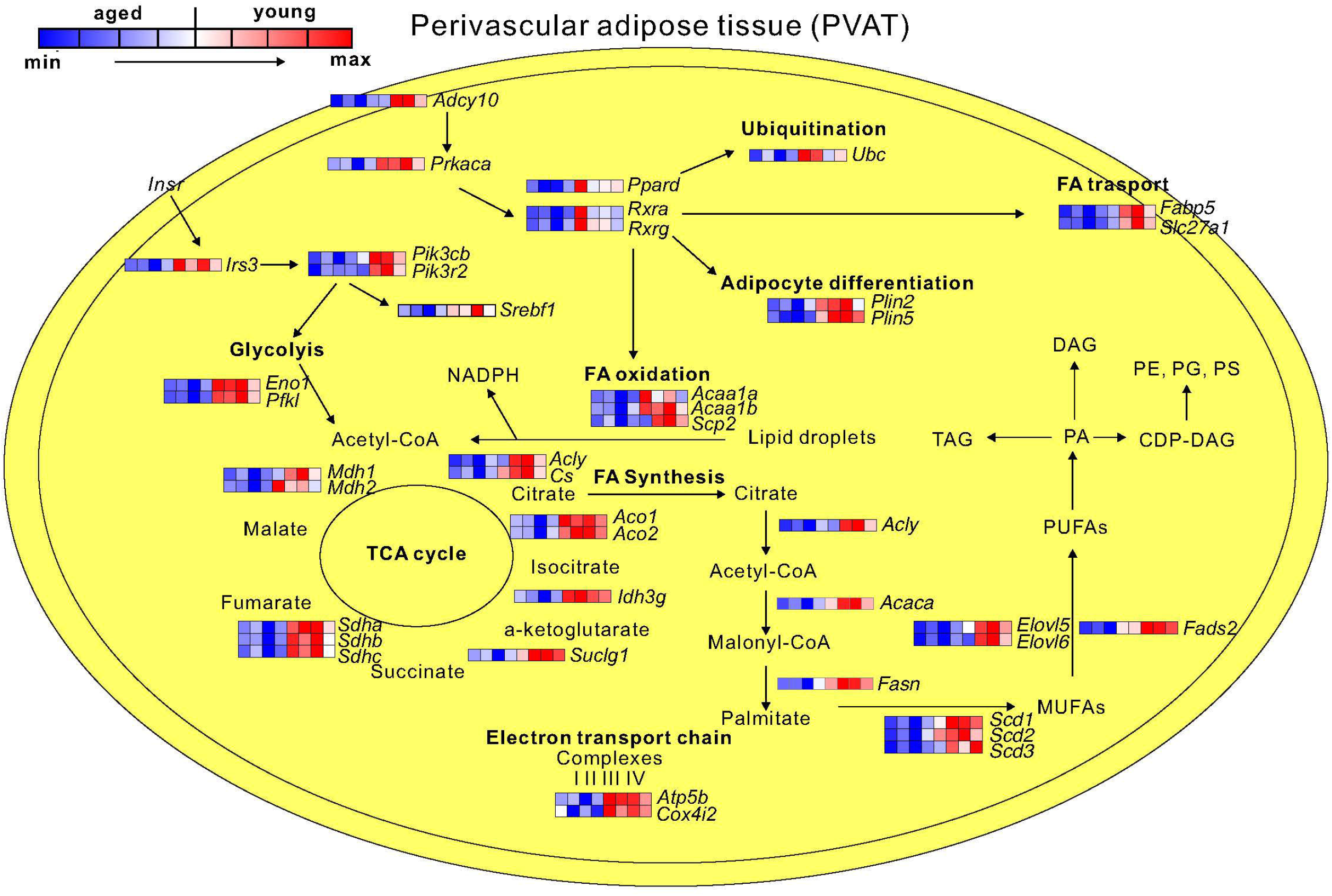
Representative schematic of downregulated genes and pathways in PVAT isolated from mesenteric arteries of 12-months old mice. By decreased expression of genes belonging to Ppar and Rcr signalling pathway, the ubiquitination, fatty acid (FA) transport, adipocyte differentiation, and FAs oxidation are decreased. In addition, the downregulated glycolysis, TCA cycle, and fatty acid biosynthesis can decrease production of perivascular fat tissue derived relaxation factors (PVATRFs) (n = 4 for young; n = 4 for 12-months old). MUFAs, monounsaturated fatty acids; PUFAs, polyunsaturated fatty acids; PA, phosphatidic acid; TAG, triacylglycerol; DAG, diacylglycerol; PE, phosphatidylethanolamine; PG, phosphatidylglycerol; PS; phosphatidylserine. Abbreviations can be found in Table S4.

## DISCUSSION

We present several novel findings. We showed that the anti-contractile effects of PVAT are impaired in mouse mesenteric arteries with increased age. Moreover, we observed altered functional role of K_V_7 (KCNQ) channels during aging. Our experiments suggest that QO58 (KCNQ channel activator) represents a novel tool for targeting KCNQ channels and possibly high blood pressure in the elderly. Furthermore, aging-related transcriptome changes of mesenteric arteries and PVAT uncover possible downstream targets of PVAT signaling pathway. Altogether, our results provide novel insights into cardiovascular events associated with aging.

Alterations in PVAT contribute to vascular dysfunction in obesity, hypertension, and cardiometabolic disease in animal models and humans (Galvez-Prieto et al., 2012) (Greenstein et al., 2009). We found that the anticontractile effects of PVAT are diminished in mouse mesenteric arteries with increased age. We conclude from these data that adipose-vascular uncoupling undergoes age-dependent changes during life span. Agabiti-Rosei et al. also found that the anticontractile effects of PVAT are abolished in mesenteric arteries from aging SAMP8 mice, which is a senescence-accelerated prone mouse model (Agabiti-Rosei et al., 2017). These findings support the notion that restoring adipose-vascular coupling could be a promising therapeutic strategy in vascular aging. We further investigated K_V_7 family of K^+^ channels as putative downstream targets of relaxing factors released by PVAT (Schleifenbaum et al., 2010) (Tsvetkov et al., 2017). Our data show that K_V_7.3-5 channel opening by flupirtine and retigabine induces relaxation in mesenteric arteries from young and old mice, making K_V_7.3-5 channels a possible therapeutic target for hypertension treatment in the elderly. Similar effects were observed for QO58, which is a novel KCNQ channel activator. In whole-cell patch-clamp and cell culture experiments, QO58 demonstrates high potency of opening KCNQ channels (for K_V_7.4 EC_50_ = 0.6 µM, for K_V_7.3/7.5 EC_50_ = 5.2 µM) (Zhang et al., 2013). Thus, QO58 might be a promising novel tool for translational research in vascular biology. Although QO58-lysine modification resulted in an improved bioavailability of the drug (Teng et al., 2016), our results in whole artery preparations argue against efficacy of QO58-lysine to be capable to open KCNQ channels in intact vascular tissue. Nevertheless, our data demonstrate that KCNQ channel induced relaxations are attenuated by all three K_V_7 channel openers tested in arteries from old mice, implying that K_V_7 function is altered in aging (Figure 2). We found that *Kcnq1,3,4,5* mRNA expression was unchanged in the arteries during aging. These RNA-seq findings were confirmed by qPCR. Thus, we concluded that impaired relaxation caused by KCNQ channels activation is not due to their changes in mRNA expression.

The RNA-Seq did not reveal additional targets or pathways. For instance, mRNA gene expressions already known to regulate KCNQ channel function (Table S5) were similar in aged mice (Figure 5). Thus, other mechanisms such as posttranslational modification (PTM) or trafficking could be responsible for age-associated KCNQ channel dysfunction. Noteworthy, PTM is a new immerging paradigm of acquired channelopathies that can occur in congestive heart failure (Curran & Mohler, 2015). Posttranslational modification of ion channels such as voltage-dependent Na channels is observed in chronic pain syndrome (Laedermann, Abriel, & Decosterd, 2015). Future studies are necessary to clarify PTM’s contribution to regulation of vascular tone in aging.

Nonetheless, RNA-Seq revealed activation of inflammatory process in old mice in mesenteric arteries. To our knowledge, this study is the first to firmly establish inflammatory transcriptome profile during different age using small resistant arteries (diameters: 150-200 μm). Our data are also consistent with the idea that inflammation is one of the key mechanisms causing vascular damage in mouse aged aorta (Gao et al., 2020). In addition, Th17 dependent immune response was activated (Table S1). In line with our previous findings, Th17 axis plays an important role in increased blood pressure (Wilck et al., 2017). Importantly, the anticontractile properties of PVAT can be restored in aging by melatonin treatment associated with decreased oxidative stress and inflammatory reaction (Agabiti-Rosei et al., 2017). Furthermore, anti-inflammatory therapy targeting the interleukin-1β innate immunity pathway in patients significantly decreased rate of recurrent cardiovascular events (Ridker et al., 2017). The participants were 60 years old on average and number needed to treat was relatively large (∼20). Since middle-aged human (38-47 years) equivalents to 1 year old mouse one could argue that anti-inflammatory interventions in human should be started earlier, in order to achieve better outcome (Flurkey, M. Currer, & Harrison, 2007). Moreover, in the later phase of aging remodeling takes place as shown by upregulated GO:0030198 and extracellular matrix organization (Table S1). Similar results were found in mouse aorta (Gao et al., 2020), suggesting that vasculature damage caused by low grade inflammation is a common process in aging.

Previous studies identified „NAD-boost” with nicotinamide mononucleotide as activator of sirtuindeacylases and a tool to reverse vascular aging (Das et al., 2018). By showing downregulation of pathway associated with energy production and therefore production of NAD and NAD precursors in mesenteric arteries of 16-months old mice (Table S2), our study contributes to the debate about the importance of NAD dependent activity of sirtuindeacylases in aging. Interestingly, PVAT transcriptional profile in our study resembles visceral fat in insulin resistance patients. For example, downregulated mitochondrial respiratory and lipid metabolic pathways were found in obese insulin-resistant subjects (Soronen et al., 2012). We observed similar pattern in PVAT genes involved into fatty acid, cholesterol and triglyceride metabolism (*Fatp2, Elovl6, Srebf1*). *Db/db* gene-deficient mice exhibited decreased expression of *Srebf1*, which was associated with impaired anti-contractile effects of PVAT (Yahagi et al., 2002) (Meijer et al., 2013). Similar results were obtained at the protein level using adipose tissue proteomic profiling in aged mice (Table S6) (Q. Yu et al., 2020). The proper function of these metabolic pathways might be essential for producing PVAT derived relaxation factors (PVATRFs), as for example PUFA. Similar metabolic pathways may control smooth muscle cell differentiation through subset of PVAT-derived stem cells (Gu et al., 2019). Thus, in addition to existing criteria such as Ca^2+^-dependence for PVATRFs, these factors could represent metabolites of fatty acids biosynthesis. Consequently, PVATRFs concentration should decrease during aging. Furthermore, PPAR pathway is compromised through peroxisome proliferator-activated receptor-γ coactivator-1 α (Ppargc1a) and lead to vascular remodeling during aging via decreased brown adipogenic differentiation in PVAT isolated from aorta (Pan et al., 2019). Our data indicate that PPAR pathway is also downregulated in PVAT surrounding mouse mesenteric arteries. However, the mechanism does unlikely involve *Ppargc1a* mRNA, since its expression was similar in aged and young mice (fold change = 0.27, padj = 0.47). Functionally distinct vessel type could explain this difference.

We studied mRNA transcript differences of several ion channels in aging vessels. Only three transcripts, namely upregulated *Cacna2d3, Kcnk2* and downregulated *P2rx5*, intersected all three data sets (12-, 16-, and 24-months old mice). We speculate that these ion channels could represent novel putative targets of arterial tone regulation. For example, auxiliary voltage-dependent calcium channel subunits delta (Cacna2d) contribute to trafficking and proper surface expression of voltage-gated calcium channels (VGCCs, Ca_V_2) (Dolphin, 2012). These channels are responsible for the P/Q current in and therefore, could be of great importance for blood pressure regulation (Andreasen et al., 2006). Interestingly, *Cacna2d3 knock out* mice exhibit reduced L-type and N-type currents in spiral ganglion neurons (Stephani et al., 2019). Thus, although the vascular phenotype of *Cacna2d3* deficient mice is not yet characterized, *Cacna2d3* arises as a novel candidate for increased blood pressure during aging. *Kcnk2* is known as TREK-1, (tandem of P domains in a weak inward-rectifier-related K^+^) channel. The channel has been implicated to play an important role in the brain vasculature (Blondeau et al., 2007). TREK-1 opening characteristics (e.g., activation by PUFAs) elevates the family as possible new targets for PVATRFs. Of note, TREK-1 deficient mice display endothelial dysfunction with decreased relaxation of mesenteric arteries (Garry et al., 2007). However, the anti-contractile effect of PVAT has remained to be studied in these mice. *P2rx5* is a purinoceptor for ATP acting as ligand-gated ion channel. Vascular smooth muscle cells from mesenteric arteries express the P2X receptors. Though, no evidence was found for a phenotype corresponding to homomeric P2X5 receptors or to heteromeric P2X1/5 receptors and the functional role of these receptors in arteries is still unclear (Lewis & Evans, 2000). Under inflammatory conditions, osteoclasts of P2rx5 gene-deficient mice have deficits in inflammasome activation and osteoclast maturation (Kim et al., 2017). However, their vascular phenotype has not yet been studied.

## EXPERIMENTAL PROCEDURES

### Mouse model

We used young (11-18 weeks old), 12-month old (50-54 weeks), 16-month old (66-69 weeks), and 24-month old (105-106 weeks) male wild-type mice C57BL/6N. Animal care followed American Physiological Society guidelines, and local authorities (Landesamt für Gesundheit und Soziales Berlin, LAGeSo) approved all protocols. Mice were housed in individually ventilated cages under standardized conditions with an artificial 12-hour dark–light cycle with free access to water and food. Animals were randomly assigned to the experimental procedures in accordance with the German legislation on protection of animals.

### Wire Myography

Mesenteric arteries were isolated after sacrifice with isoflurane anesthesia, as previously described (Tsvetkov et al., 2017). Then blood vessels were quickly transferred to cold (4°C), oxygenated (95% O_2_ / 5% CO_2_) physiological salt solution (PSS) containing (in mmol/L) 119 NaCl, 4.7 KCl, 1.2 KH_2_PO_4_, 25 NaHCO_3_, 1.2 Mg_2_SO_4_, 11.1 glucose, 1.6 CaCl_2_). We dissected the vessels into 2 mm rings whereby perivascular fat and connective tissue were either intact ((+) PVAT) or removed ((–) PVAT rings). Each ring was placed between two stainless steel wires (diameter 0.0394 mm) in a 5-ml organ bath of a Mulvany Small Vessel Myograph (DMT 610 M; Danish Myo Technology, Denmark). The organ bath was filled with PSS. Continuously oxygenated bath solution with a gas mixture of 95% O_2_ and 5% CO_2_ were kept at 37 °C (pH 7.4). To obtain the passive diameter of the vessel at 100 mm Hg a DMT normalization procedure was performed. The mesenteric artery rings were placed under a tension equivalent to that generated at 0.9 times the diameter of the vessel at 100 mm Hg by stepwise distending the vessel using LabChart DMT Normalization module. The software Chart5 (AD Instruments Ltd. Spechbach, Germany) was used for data acquisition and display. After 60 minutes incubation arteries were pre-contracted either with isotonic external 60 mM KCl or 1-3 µM phenylephrine (PE), or methoxamine (ME) until a stable resting tension was acquired. The composition of 60 mM KCl (in mmol/L) was 63.7 NaCl, 60 KCl, 1.2 KH_2_PO_4_, 25 NaHCO_3_, 1.2 Mg_2_SO_4_, 11.1 glucose, and 1.6 CaCl_2_. Drugs were added to the bath solution if not indicated otherwise. Tension is expressed as a percentage of the steady-state tension (100%) obtained with isotonic external 60 mM KCl or agonist (e.g. PE, ME).

### Quantitative real-time PCR

Total RNA was isolated from young, 12-, 16-, and 24 months old mice mesenteric arteries (first branches) by using the RNeasy RNA isolation kit (Qiagen, Germantown, MD) according to the manufacturer’s instruction. Isolated RNA concentration was measured and RNA quality was tested by NanoDrop-1000 spectrophotometer (Thermo Fisher Scientific, Vernon Hills, IL). Two micrograms of RNA were used for cDNA transcription (Applied Biosystems, Foster City, CA). Experiments were run on an Applied Biosystems 7500 Fast Real-Time PCR System (Life Technologies Corporation, Carlsbad, CA, USA). Primers were designed using Primer 3 software on different exons to exclude any DNA contamination. Specificity of amplified products was validated in silico (blast) and empirically with gel electrophoresis and analysis of melt curves. Primers were synthesized by BioTez (Berlin, Germany); the sequences are provided below. The cycling conditions were the following: initial activation at 95°C for 10 min, followed by 40 cycles at 95°C for 15 s and 60°C for 1 min. Samples and negative controls were run in parallel. Quantitative analysis of target mRNA expression was performed with quantitative real-time PCR using the relative standard curve method. The expression level of the target genes was normalized by the expression of *18s*. Under our experimental conditions, expression of *18s* as a reference gene did not differ between young and old mice tissues. The fold change in gene expression between young and old mice was calculated using 2 ΔΔCt method. The following primers were used:

*18s*: F: 5′-ACATCCAAGGAAGGCAGCAG-3′;

R: 5′-TTTTCGTCACTACCTCCCCG-3′.

*Kcnq1*: F: 5′-AGCAGTATGCCGCTCTGG-3′;

R: 5′-AGATGCCCACGTACTTGCTG-3′.

*Kcnq3*: F:5′-CAGTATTCGGCCGGACATCT-3′;

R:5′-GAGACTGCTGGGATGGGTAG-3′.

*Kcnq4*: F:5′-CACTTTGAGAAGCGCAGGAT-3′;

R:5′-CCAGGTGGCTGTCAAATAGG-3′.

*Kcnq5*: F:5′-CCTCACTACGGCTCAAGAGT-3′;

R:5′-TTAAGTGGTGGGGTGAGGTC-3′.

### RNA Sequencing

Following Agilent 2100 bioanalyzer quality control, RNA-seq was performed using Illumina Genome Analyzer Novaseq 6000 platform. NEB Next® Ultra™ RNA Library Prep Kit was used for library preparation. Sequence quality estimations, GC content, nucleotide distribution, and read duplication levels were determined for the samples using fastp-0.12.2 software. The reads were mapped to the reference mouse genome (ensembl_mus_musculus_grcm38_p6_gca_000001635_8). HISAT2 was selected to map the filtered sequenced reads to the reference genome. The uniquely mapped read data output was processed using custom scripts in R software (version 3.5.1), then normalized using the FeatureCounts package v1.5.0-p3 version. Differential expression analysis was performed using the DESeq2 R package version v1.20.0 (Anders & Huber, 2010). We used clusterProfiler for enrichment analysis, including GO Enrichment, DO Enrichment, KEGG and Reactome database Enrichment (G. Yu, Wang, Han, & He, 2012). Heat map was generated based on fragments per kilobase per million mapped fragments (FPKM) values using Morpheus software (https://software.broadinstitute.org/morpheus).

### Materials

All salts and other chemicals were purchased from Sigma-Aldrich (Germany) or Merck (Germany). Using DMSO or PSS, drugs were freshly dissolved on the day of each experiment accordingly to the material sheet. Maximal DMSO concentration after application did not exceed 0.5%. Following concentration of drugs were used: phenylephrine (Sigma Aldrich) and methoxamine (Sigma Aldrich) ranged from 0.01 to 100 µM; retigabine (Valeant Research North America), flupirtine (Tocris), QO58 (Tocris), QO58-lysine from 0.01 to 30 µM; 3 µM XE991 (Tocris).

### Statistics

Data present mean ± SEM. We calculated EC_50_ values using a Hill equation: T = (B_0_ – Be) / (1 + ([D]/EC_50_)^n^) + Be, where T is the tension in response to the drug (D); Be is the maximum response induced by the drug; B_0_, is a constant; EC_50_ is the concentration of the drug that elicits a half-maximal response.

For curve fittings using non-linear regression GraphPad 8.0.1 (Software, La Jolla California USA) software was used. Statistical significance was determined by Mann-Whitney test or nonparametric ANOVA (Kruskal-Wallis test). Extra sum-of-squares F test was performed for comparison of concentration-response curves. *P* values < 0.05 were considered statistically significant. n represents the number of arteries tested. Figures were made using CorelDRAW Graphics Suite 2020 (Ottawa, Canada).

## ACKNOWLEDGMENTS

We thank Fan Zhang, Kewei Wang, Hi-lin Zhang for providing QO58-lysine and Zhihuang Zheng for assistance with quantitative real[time PCR.

## AUTHOR CONTRIBUTIONS

Y.W., F.Y., A.S., M.K, F.L., M.G., and D.T. were responsible for data collection, analysis, and interpretation. Y.W. and D.T. drafted the manuscript. All authors have approved the final version of the manuscript and agreed to be accountable for all aspects of the work. All persons designated as authors qualify for authorship, and all those who qualify for authorship are listed.

## DATA AVAILABILITY STATEMENT

All data used in this study will be made available.

## REFERENCES

Agabiti-Rosei, C., Favero, G., De Ciuceis, C., Rossini, C., Porteri, E., Rodella, L. F., Rezzani, R. (2017). Effect of long-term treatment with melatonin on vascular markers of oxidative stress/inflammation and on the anticontractile activity of perivascular fat in aging mice. Hypertens Res, 40(1), 41–50. doi:10.1038/hr.2016.103

Anders, S., & Huber, W. (2010). Differential expression analysis for sequence count data. Genome Biol, 11(10), R106. doi:10.1186/gb-2010-11-10-r106

Andreasen, D., Friis, U. G., Uhrenholt, T. R., Jensen, B. L., Skott, O., & Hansen, P. B. (2006). Coexpression of voltage-dependent calcium channels Cav1.2, 2.1a, and 2.1b in vascular myocytes. Hypertension, 47(4), 735–741. doi:10.1161/01.HYP.0000203160.80972.47

Barrese, V., Stott, J. B., & Greenwood, I. A. (2018). KCNQ-Encoded Potassium Channels as Therapeutic Targets. Annu Rev Pharmacol Toxicol, 58, 625–648. doi:10.1146/annurev-pharmtox-010617-052912

Blondeau, N., Petrault, O., Manta, S., Giordanengo, V., Gounon, P., Bordet, R., Heurteaux, C. (2007). Polyunsaturated fatty acids are cerebral vasodilators via the TREK-1 potassium channel. Circ Res, 101(2), 176–184. doi:10.1161/CIRCRESAHA.107.154443

Collaborators, G. B. D. R. F. (2020). Global burden of 87 risk factors in 204 countries and territories, 1990-2019: a systematic analysis for the Global Burden of Disease Study 2019. Lancet, 396(10258), 1223–1249. doi:10.1016/S0140-6736(20)30752-2

Curran, J., & Mohler, P. J. (2015). Alternative paradigms for ion channelopathies: disorders of ion channel membrane trafficking and posttranslational modification. Annu Rev Physiol, 77, 505–524. doi:10.1146/annurev-physiol-021014-071838

Das, A., Huang, G. X., Bonkowski, M. S., Longchamp, A., Li, C., Schultz, M. B., Sinclair, D. A. (2018). Impairment of an Endothelial NAD(+)-H2S Signaling Network Is a Reversible Cause of Vascular Aging. Cell, 173(1), 74–89 e20. doi:10.1016/j.cell.2018.02.008

Dolphin, A. C. (2012). Calcium channel auxiliary alpha2delta and beta subunits: trafficking and one step beyond. Nat Rev Neurosci, 13(8), 542–555. doi:10.1038/nrn3311

DuPont, J. J., McCurley, A., Davel, A. P., McCarthy, J., Bender, S. B., Hong, K., Jaffe, I. Z. (2016). Vascular mineralocorticoid receptor regulates microRNA-155 to promote vasoconstriction and rising blood pressure with aging. JCI Insight, 1(14), e88942. doi:10.1172/jci.insight.88942

Essin, K., Welling, A., Hofmann, F., Luft, F. C., Gollasch, M., & Moosmang, S. (2007). Indirect coupling between Cav1.2 channels and ryanodine receptors to generate Ca2+ sparks in murine arterial smooth muscle cells. J Physiol, 584(Pt 1), 205–219. doi:10.1113/jphysiol.2007.138982

European.Medicines.Agency. (2018). Withdrawal of pain medicine flupirtine endorsed.

FDA. (2013). FDA determines 2013 labeling adequate to manage risk of retinal abnormalities, potential vision loss, and skin discoloration with anti-seizure drug Potiga (ezogabine). FDA Drug Safety Communication.

Flurkey, K. M., Currer, J., & Harrison, D. E. (2007). Chapter 20 - Mouse Models in Aging Research. In J. G. Fox, M. T. Davisson, F. W. Quimby, S. W. Barthold, C. E. Newcomer, & A. L. Smith (Eds.), The Mouse in Biomedical Research (Second Edition) (pp. 637–672). Burlington: Academic Press.

Galvez-Prieto, B., Somoza, B., Gil-Ortega, M., Garcia-Prieto, C. F., de Las Heras, A.I., Gonzalez, M. C., Fernandez-Alfonso, M. S. (2012). Anticontractile Effect of Perivascular Adipose Tissue and Leptin are Reduced in Hypertension. Front Pharmacol, 3, 103. doi:10.3389/fphar.2012.00103

Gao, P., Gao, P., Choi, M., Chegireddy, K., Slivano, O. J., Zhao, J., Long, X. (2020). Transcriptome analysis of mouse aortae reveals multiple novel pathways regulated by aging. Aging (Albany NY), 12(15), 15603–15623. doi:10.18632/aging.103652

Garry, A., Fromy, B., Blondeau, N., Henrion, D., Brau, F., Gounon, P., Saumet, J. L. (2007). Altered acetylcholine, bradykinin and cutaneous pressure-induced vasodilation in mice lacking the TREK1 potassium channel: the endothelial link. EMBO Rep, 8(4), 354–359. doi:10.1038/sj.embor.7400916

Gollasch, M. (2017). Adipose-Vascular Coupling and Potential Therapeutics. Annu Rev Pharmacol Toxicol, 57, 417–436. doi:10.1146/annurev-pharmtox-010716-104542

Greenstein, A. S., Khavandi, K., Withers, S. B., Sonoyama, K., Clancy, O., Jeziorska, M., Heagerty, A. M. (2009). Local inflammation and hypoxia abolish the protective anticontractile properties of perivascular fat in obese patients. Circulation, 119(12), 1661–1670. doi:10.1161/CIRCULATIONAHA.108.821181

Gu, W., Nowak, W. N., Xie, Y., Le Bras, A., Hu, Y., Deng, J., Xu, Q. (2019). Single-Cell RNA-Sequencing and Metabolomics Analyses Reveal the Contribution of Perivascular Adipose Tissue Stem Cells to Vascular Remodeling. Arterioscler Thromb Vasc Biol, 39(10), 2049–2066. doi:10.1161/ATVBAHA.119.312732

Jepps, T. A., Chadha, P. S., Davis, A. J., Harhun, M. I., Cockerill, G. W., Olesen, S. P., Greenwood, I. A. (2011). Downregulation of Kv7.4 channel activity in primary and secondary hypertension. Circulation, 124(5), 602–611. doi:10.1161/CIRCULATIONAHA.111.032136

Jia, C., Qi, J., Zhang, F., Mi, Y., Zhang, X., Chen, X., Zhang, H. (2011). Activation of KCNQ2/3 potassium channels by novel pyrazolo[1,5-a]pyrimidin-7(4H)-one derivatives. Pharmacology, 87(5-6), 297–310. doi:10.1159/000327384

Kim, H., Walsh, M. C., Takegahara, N., Middleton, S. A., Shin, H. I., Kim, J., & Choi, Y. (2017). The purinergic receptor P2X5 regulates inflammasome activity and hyper-multinucleation of murine osteoclasts. Sci Rep, 7(1), 196. doi:10.1038/s41598-017-00139-2

Laedermann, C. J., Abriel, H., & Decosterd, I. (2015). Post-translational modifications of voltage-gated sodium channels in chronic pain syndromes. Front Pharmacol, 6, 263. doi:10.3389/fphar.2015.00263

Lewis, C. J., & Evans, R. J. (2000). Lack of run-down of smooth muscle P2X receptor currents recorded with the amphotericin permeabilized patch technique, physiological and pharmacological characterization of the properties of mesenteric artery P2X receptor ion channels. Br J Pharmacol, 131(8), 1659–1666. doi:10.1038/sj.bjp.0703744

Lohn, M., Dubrovska, G., Lauterbach, B., Luft, F. C., Gollasch, M., & Sharma, A. M. (2002). Periadventitial fat releases a vascular relaxing factor. FASEB J, 16(9), 1057–1063. doi:10.1096/fj.02-0024com

Mani, B. K., Robakowski, C., Brueggemann, L. I., Cribbs, L. L., Tripathi, A., Majetschak, M., & Byron, K. L. (2016). Kv7.5 Potassium Channel Subunits Are the Primary Targets for PKA-Dependent Enhancement of Vascular Smooth Muscle Kv7 Currents. Mol Pharmacol, 89(3), 323–334. doi:10.1124/mol.115.101758

Meijer, R. I., Bakker, W., Alta, C. L., Sipkema, P., Yudkin, J. S., Viollet, B., Eringa, E. C. (2013). Perivascular adipose tissue control of insulin-induced vasoreactivity in muscle is impaired in db/db mice. Diabetes, 62(2), 590–598. doi:10.2337/db11-1603

Morales-Cano, D., Moreno, L., Barreira, B., Pandolfi, R., Chamorro, V., Jimenez, R., Cogolludo, A. (2015). Kv7 channels critically determine coronary artery reactivity: left-right differences and down-regulation by hyperglycaemia. Cardiovasc Res, 106(1), 98–108. doi:10.1093/cvr/cvv020

North, B. J., & Sinclair, D. A. (2012). The intersection between aging and cardiovascular disease. Circ Res, 110(8), 1097–1108. doi:10.1161/CIRCRESAHA.111.246876

Pan, X. X., Ruan, C. C., Liu, X. Y., Kong, L. R., Ma, Y., Wu, Q. H., Gao, P. J. (2019). Perivascular adipose tissue-derived stromal cells contribute to vascular remodeling during aging. Aging Cell, 18(4), e12969. doi:10.1111/acel.12969

Ridker, P. M., Everett, B. M., Thuren, T., MacFadyen, J. G., Chang, W. H., Ballantyne, C., Group, C. T. (2017). Antiinflammatory Therapy with Canakinumab for Atherosclerotic Disease. N Engl J Med, 377(12), 1119–1131. doi:10.1056/NEJMoa1707914

Schleifenbaum, J., Kohn, C., Voblova, N., Dubrovska, G., Zavarirskaya, O., Gloe, T., Gollasch, M. (2010). Systemic peripheral artery relaxation by KCNQ channel openers and hydrogen sulfide. J Hypertens, 28(9), 1875–1882. doi:10.1097/HJH.0b013e32833c20d5

Soronen, J., Laurila, P. P., Naukkarinen, J., Surakka, I., Ripatti, S., Jauhiainen, M., Yki-Jarvinen, H. (2012). Adipose tissue gene expression analysis reveals changes in inflammatory, mitochondrial respiratory and lipid metabolic pathways in obese insulin-resistant subjects. BMC Med Genomics, 5, 9. doi:10.1186/1755-8794-5-9

Stephani, F., Scheuer, V., Eckrich, T., Blum, K., Wang, W., Obermair, G. J., & Engel, J. (2019). Deletion of the Ca(2+) Channel Subunit alpha2delta3 Differentially Affects Cav2.1 and Cav2.2 Currents in Cultured Spiral Ganglion Neurons Before and After the Onset of Hearing. Front Cell Neurosci, 13, 278. doi:10.3389/fncel.2019.00278

Tabula Muris, C. (2020). A single-cell transcriptomic atlas characterizes ageing tissues in the mouse. Nature, 583(7817), 590–595. doi:10.1038/s41586-020-2496-1

Teng, B. C., Song, Y., Zhang, F., Ma, T. Y., Qi, J. L., Zhang, H. L., Wang, K. (2016). Activation of neuronal Kv7/KCNQ/M-channels by the opener QO58-lysine and its anti-nociceptive effects on inflammatory pain in rodents. Acta Pharmacol Sin, 37(8), 1054–1062. doi:10.1038/aps.2016.33

Tsvetkov, D., Kassmann, M., Tano, J. Y., Chen, L., Schleifenbaum, J., Voelkl, J., Gollasch, M. (2017). Do KV 7.1 channels contribute to control of arterial vascular tone? Br J Pharmacol, 174(2), 150–162. doi:10.1111/bph.13665

Verlohren, S., Dubrovska, G., Tsang, S. Y., Essin, K., Luft, F. C., Huang, Y., & Gollasch, M. (2004). Visceral periadventitial adipose tissue regulates arterial tone of mesenteric arteries. Hypertension, 44(3), 271–276. doi:10.1161/01.HYP.0000140058.28994.ec

Vollset, S. E., Goren, E., Yuan, C. W., Cao, J., Smith, A. E., Hsiao, T., Murray, C. J. L. (2020). Fertility, mortality, migration, and population scenarios for 195 countries and territories from 2017 to 2100: a forecasting analysis for the Global Burden of Disease Study. Lancet, 396(10258), 1285–1306. doi:10.1016/S0140-6736(20)30677-2

Wilck, N., Matus, M. G., Kearney, S. M., Olesen, S. W., Forslund, K., Bartolomaeus, H., Muller, D. N. (2017). Salt-responsive gut commensal modulates TH17 axis and disease. Nature, 551(7682), 585–589. doi:10.1038/nature24628

Yahagi, N., Shimano, H., Hasty, A. H., Matsuzaka, T., Ide, T., Yoshikawa, T., Yamada, N. (2002). Absence of sterol regulatory element-binding protein-1 (SREBP-1) ameliorates fatty livers but not obesity or insulin resistance in Lep(ob)/Lep(ob) mice. J Biol Chem, 277(22), 19353–19357. doi:10.1074/jbc.M201584200

Yu, G., Wang, L. G., Han, Y., & He, Q. Y. (2012). clusterProfiler: an R package for comparing biological themes among gene clusters. OMICS, 16(5), 284–287. doi:10.1089/omi.2011.0118

Yu, Q., Xiao, H., Jedrychowski, M. P., Schweppe, D. K., Navarrete-Perea, J., Knott, J., Gygi, S. P. (2020). Sample multiplexing for targeted pathway proteomics in aging mice. Proc Natl Acad Sci U S A, 117(18), 9723–9732. doi:10.1073/pnas.1919410117

Zhang, F., Mi, Y., Qi, J. L., Li, J. W., Si, M., Guan, B. C., Zhang, H. L. (2013). Modulation of K(v)7 potassium channels by a novel opener pyrazolo[1,5-a]pyrimidin-7(4H)-one compound QO-58. Br J Pharmacol, 168(4), 1030–1042. doi:10.1111/j.1476-5381.2012.02232.x

