## Supplemental Figure 1, Supplemental Table1-5, for "Aging affects K_V_7 channels and perivascular-adipose tissue-mediated vascular tone"

**DATA SUPPLEMENT**

**Figure S1**

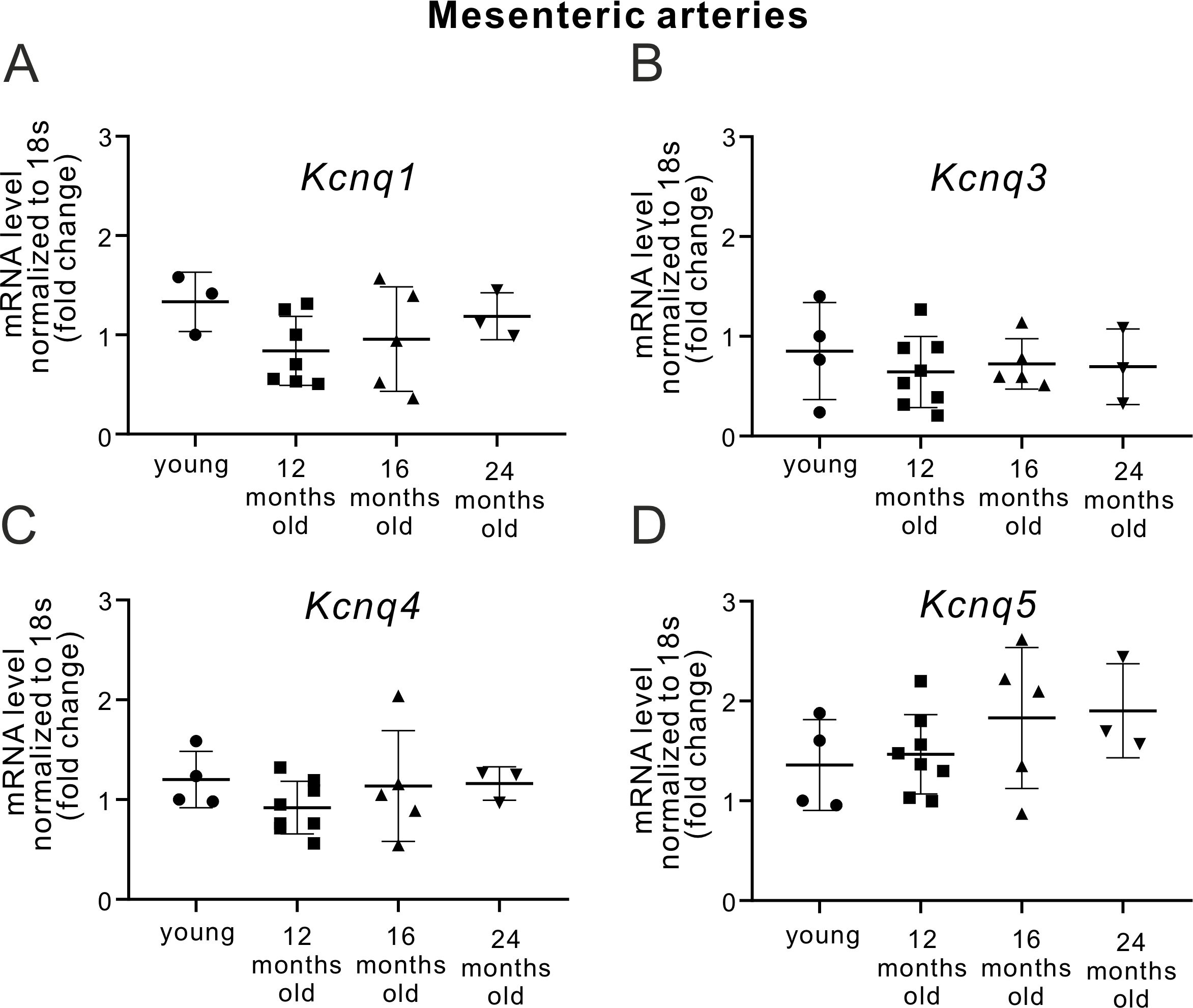

**Figure S1.** Relative expression of KCNQ *1, 3, 4, 5* channels at mRNA levels in (–) PVAT mesenteric arteries from young and aged mice normalized to *18s*.

(A) Relative mRNA levels for *Kcnq1* (n = 3 for young; n = 7 for 12-months old; n = 5 for 16-months old; n = 3 for 24 months old mice).

(B) Relative mRNA levels for *Kcnq3* (n = 4 for young; n = 8 for 12-months old; n = 5 for 16-months old; n = 3 for 24-months old mice).

(C) Relative mRNA levels for *Kcnq4* (n = 4 for young; n = 8 for 12-months old; n = 5, for 16-months old; n = 3 for 24-months old mice).

(D) Relative mRNA levels for *Kcnq5* (n = 4 for young; n = 8 for 12-months old; n = 5 for 16-months old; n = 3 for 24-months old mice). ns, P > 0.05, Kruskal–Wallis one-way analysis of variance. Data are mean and SD.

**Table S1. Top upregulated GO Terms and KEGG Pathways in PVAT isolated from 12-month old mice**

| **GO Biological Process** | | | |
| --- | --- | --- | --- |
| **id** | **Terms** | **p. value** | **Adj.p.value** |
| GO:0002250 | adaptive immune response | 1.18E-90 | 6.11E-87 |
| GO:0051249 | regulation of lymphocyte activation | 1.52E-75 | 3.91E-72 |
| GO:0002764 | immune response-regulating signaling pathway | 5.80E-75 | 9.98E-72 |
| GO:0002757 | immune response-activating signal transduction | 4.56E-72 | 5.89E-69 |
| GO:0050867 | positive regulation of cell activation | 7.06E-71 | 7.30E-68 |
| GO:0098542 | defense response to other organism | 9.32E-71 | 8.02E-68 |
| GO:0002253 | activation of immune response | 2.83E-70 | 2.09E-67 |
| GO:0002696 | positive regulation of leukocyte activation | 4.29E-69 | 2.77E-66 |
| GO:0042113 | B cell activation | 1.70E-68 | 9.78E-66 |
| GO:0002768 | immune response-regulating cell surface receptor signaling pathway | 6.12E-67 | 3.16E-64 |
| GO:0051251 | positive regulation of lymphocyte activation | 1.54E-66 | 7.21E-64 |
| GO:0002429 | immune response-activating cell surface receptor signaling pathway | 1.36E-64 | 5.85E-62 |
| GO:0050851 | antigen receptor-mediated signaling pathway | 5.76E-64 | 2.29E-61 |
| GO:0002460 | adaptive immune response based on somatic recombination of immune receptors built from immunoglobulin superfamily domains | 1.23E-63 | 4.53E-61 |
| GO:0002443 | leukocyte mediated immunity | 2.14E-60 | 7.36E-58 |
| GO:0002449 | lymphocyte mediated immunity | 1.72E-58 | 5.56E-56 |
| GO:0050864 | regulation of B cell activation | 9.82E-57 | 2.98E-54 |
| GO:0050853 | B cell receptor signaling pathway | 6.90E-56 | 1.98E-53 |
| GO:0042110 | T cell activation | 1.30E-51 | 3.53E-49 |
| GO:0002250 | adaptive immune response | 1.18E-90 | 6.11E-87 |
| **KEGG** | | | |
| **id** | **Terms** | **p. value** | **Adj.p.value** |
| mmu04060 | Cytokine-cytokine receptor interaction | 1.35E-24 | 4.06E-22 |
| mmu05340 | Primary immunodeficiency | 7.28E-23 | 1.09E-20 |
| mmu04640 | Hematopoietic cell lineage | 3.45E-22 | 3.45E-20 |
| mmu04672 | Intestinal immune network for IgA production | 2.17E-17 | 1.62E-15 |
| mmu04380 | Osteoclast differentiation | 1.00E-16 | 6.02E-15 |
| mmu04658 | Th1 and Th2 cell differentiation | 1.48E-16 | 7.42E-15 |
| mmu05321 | Inflammatory bowel disease (IBD) | 2.26E-16 | 9.47E-15 |
| mmu04064 | NF-kappa B signaling pathway | 2.52E-16 | 9.47E-15 |
| mmu04650 | Natural killer cell mediated cytotoxicity | 3.40E-16 | 1.13E-14 |
| mmu05330 | Allograft rejection | 1.17E-15 | 3.51E-14 |
| mmu04660 | T cell receptor signaling pathway | 1.47E-15 | 4.02E-14 |
| mmu04514 | Cell adhesion molecules (CAMs) | 1.82E-15 | 4.56E-14 |
| mmu04659 | Th17 cell differentiation | 2.98E-15 | 6.89E-14 |
| mmu04662 | B cell receptor signaling pathway | 3.09E-14 | 6.62E-13 |
| mmu05332 | Graft-versus-host disease | 4.92E-14 | 9.85E-13 |
| mmu04061 | Viral protein interaction with cytokine and cytokine receptor | 6.91E-14 | 1.30E-12 |
| mmu05168 | Herpes simplex virus 1 infection | 6.48E-13 | 1.14E-11 |
| mmu04940 | Type I diabetes mellitus | 2.12E-12 | 3.53E-11 |
| mmu05140 | Leishmaniasis | 3.59E-12 | 5.67E-11 |
| mmu04060 | Cytokine-cytokine receptor interaction | 1.35E-24 | 4.06E-22 |

**Table S2. Top downregulated GO Terms and KEGG Pathways in PVAT isolated from 12-month old mice**

| **GO Biological Process** | | | |
| --- | --- | --- | --- |
| **id** | **Terms** | **p. value** | **Adj.p.value** |
| GO:0006091 | generation of precursor metabolites and energy | 7.32E-47 | 3.48E-43 |
| GO:0051186 | cofactor metabolic process | 4.73E-41 | 1.12E-37 |
| GO:0022900 | electron transport chain | 2.51E-39 | 3.98E-36 |
| GO:0045333 | cellular respiration | 9.33E-39 | 1.11E-35 |
| GO:0046034 | ATP metabolic process | 1.84E-36 | 1.75E-33 |
| GO:0032787 | monocarboxylic acid metabolic process | 7.77E-36 | 6.15E-33 |
| GO:0015980 | energy derivation by oxidation of organic compounds | 9.13E-35 | 6.20E-32 |
| GO:0009161 | ribonucleoside monophosphate metabolic process | 4.17E-34 | 2.48E-31 |
| GO:0009167 | purine ribonucleoside monophosphate metabolic process | 6.48E-34 | 3.42E-31 |
| GO:0006119 | oxidative phosphorylation | 7.45E-34 | 3.54E-31 |
| GO:0009199 | ribonucleoside triphosphate metabolic process | 8.44E-34 | 3.65E-31 |
| GO:0009126 | purine nucleoside monophosphate metabolic process | 9.63E-34 | 3.81E-31 |
| GO:0009205 | purine ribonucleoside triphosphate metabolic process | 2.11E-33 | 7.73E-31 |
| GO:0009144 | purine nucleoside triphosphate metabolic process | 3.64E-33 | 1.23E-30 |
| GO:0009123 | nucleoside monophosphate metabolic process | 6.08E-33 | 1.93E-30 |
| GO:0022904 | respiratory electron transport chain | 1.38E-32 | 4.11E-30 |
| GO:0009141 | nucleoside triphosphate metabolic process | 4.63E-32 | 1.29E-29 |
| GO:0007005 | mitochondrion organization | 4.26E-31 | 1.12E-28 |
| GO:0042773 | ATP synthesis coupled electron transport | 1.32E-30 | 3.30E-28 |
| GO:0042775 | mitochondrial ATP synthesis coupled electron transport | 2.14E-30 | 5.08E-28 |
| **KEGG** | | | |
| **id** | **Terms** | **p. value** | **Adj.p.value** |
| mmu05012 | Parkinson disease | 1.23E-41 | 3.79E-39 |
| mmu00190 | Oxidative phosphorylation | 7.22E-41 | 1.11E-38 |
| mmu05016 | Huntington disease | 3.95E-33 | 3.64E-31 |
| mmu04932 | Non-alcoholic fatty liver disease (NAFLD) | 4.74E-33 | 3.64E-31 |
| mmu04714 | Thermogenesis | 1.07E-30 | 6.59E-29 |
| mmu01200 | Carbon metabolism | 9.08E-19 | 4.64E-17 |
| mmu01212 | Fatty acid metabolism | 3.19E-18 | 1.40E-16 |
| mmu04723 | Retrograde endocannabinoid signaling | 8.22E-15 | 3.16E-13 |
| mmu04146 | Peroxisome | 2.66E-11 | 9.07E-10 |
| mmu00620 | Pyruvate metabolism | 3.06E-11 | 9.40E-10 |
| mmu03320 | PPAR signaling pathway | 3.68E-11 | 1.03E-09 |
| mmu00020 | Citrate cycle (TCA cycle) | 2.73E-10 | 6.99E-09 |
| mmu00280 | Valine, leucine and isoleucine degradation | 4.99E-10 | 1.18E-08 |
| mmu00010 | Glycolysis / Gluconeogenesis | 2.95E-09 | 6.34E-08 |
| mmu04260 | Cardiac muscle contraction | 3.10E-09 | 6.34E-08 |
| mmu01040 | Biosynthesis of unsaturated fatty acids | 7.96E-09 | 1.53E-07 |
| mmu01230 | Biosynthesis of amino acids | 4.98E-08 | 8.99E-07 |
| mmu00062 | Fatty acid elongation | 6.78E-08 | 1.16E-06 |
| mmu00640 | Propanoate metabolism | 3.46E-07 | 5.58E-06 |
| mmu00900 | Terpenoid backbone biosynthesis | 5.77E-07 | 8.86E-06 |

**Table S3. Abbreviations used in Figure 5**

| **Gene Symbol** | **Gene Description** |
| --- | --- |
| *Fgf14* | Fibroblast growth factor 14 |
| *Kcne1l* | Potassium voltage-gated channel subfamily E regulatory beta subunit 5 |
| *Kcne2* | Potassium voltage-gated channel subfamily E Isk-related subfamily 2 |
| *Kcne3* | Potassium voltage-gated channel subfamily E member 3 |
| *Kcne4* | Potassium voltage-gated channel subfamily E member 4 |
| *Nedd4l* | Neural precursor cell expressed, developmentally down-regulated 4-like, E3 ubiquitin protein ligase |
| *Plcb3* | phospholipase C, beta 3 |
| *Plcb4* | phospholipase C, beta 4 |
| *Plcd4* | phospholipase C, delta 4 |
| *Plcxd1* | PI-PLC X domain-containing protein 1 |
| *Plcxd2* | PI-PLC X domain-containing protein 2 |
| *Plcxd3* | Phosphatidylinositol-specific phospholipase C, X domain containing 3 |
| *Prkaca* | protein kinase, cAMP dependent, catalytic, alpha |
| *Sgk1* | Serine/threonine-protein kinase Sgk1 |
| *Slc5a3* | Sodium/myo-inositol cotransporter |
| *Cyp1a1* | Cytochrome P450 1A1 |
| *Wisp2* | WNT1 inducible signaling pathway protein 2 |
| *Pon1* | Paraoxonase 1 |
| *8430408G22* | Protein DEPP1 |
| *Apoc1* | Apolipoprotein C-I Truncated apolipoprotein C-I |
| *Csprs* | Component of Sp100-rs |
| *Gm15433* | predicted pseudogene 15433 |
| *Cd22* | CD22 molecule |
| *Ms4ab* | Membrane-spanning 4-domains, subfamily A, member 4B |
| *Ms4a1* | Membrane-spanning 4-domains, subfamily A, member 1 |

**Table S4. Abbreviations used in Figure 6**

| **Gene Symbol** | **Gene Description** |
| --- | --- |
| *Adcy10* | Adenylate cyclase type 10 |
| *Prkaca* | Protein kinase, cAMP dependent, catalytic, alpha |
| *Ppard* | Peroxisome proliferator-activated receptor delta |
| *Rxra* | Retinoid X receptor alpha |
| *Rxrg* | Retinoic acid receptor RXR-gamma |
| *Ubc* | Ubiquitin C |
| *Fabp5* | Fatty acid-binding protein |
| *Plin2* | Perilipin-2 |
| *Plin5* | Perilipin-5 |
| *Acaa1a* | Acetyl-Coenzyme A acyltransferase 1A |
| *Acaa1b* | Acetyl-Coenzyme A acyltransferase 1B |
| *Scp2* | Sterol carrier protein 2 |
| *Insr* | Insulin receptor |
| *Irs3* | Insulin receptor substrate 3 |
| *Pik3cb* | Phosphatidylinositol 4,5-bisphosphate 3-kinase catalytic subunit beta isoform |
| *Pik3r2* | Phosphatidylinositol 3-kinase regulatory subunit beta |
| *Srebf1* | Sterol regulatory element-binding protein 1 |
| *Eno1* | Alpha-enolase |
| *Pfkl* | ATP-dependent 6-phosphofructokinase |
| *Acly* | ATP citrate lyase |
| *Cs* | Citrate synthase |
| *Aco1* | Aconitase 1 |
| *Aco2* | Aconitase 2 |
| *Idh3g* | Isocitrate dehydrogenase 3 (NAD+), gamma |
| *Suclg1* | Succinate-CoA ligase [ADP/GDP-forming] subunit alpha |
| *Sdha* | Succinate dehydrogenase complex, subunit A |
| *Sdhb* | Succinate dehydrogenase complex, subunit B |
| *Sdhc* | Succinate dehydrogenase complex, subunit C |
| *Mdh1* | Malate dehydrogenase 1 |
| *Mdh2* | Malate dehydrogenase 2 |
| *Acaca* | Acetyl-Coenzyme A carboxylase alpha |
| *Fasn* | Fatty acid synthase |
| *Atp5b* | ATP synthase subunit beta |
| *Cox4i2* | Cytochrome c oxidase subunit 4 isoform 2 |
| *Scd1* | Acyl-CoA desaturase 1 |
| *Scd2* | Acyl-CoA desaturase 2 |
| *Scd3* | Acyl-CoA desaturase 3 |
| *Fads2* | Fatty acid desaturase 2 |
| *Elovl5* | Elongation of very long-chain fatty acid protein 5 |
| *Elovl6* | Elongation of very long-chain fatty acid protein 6 |

**Table S5. Candidates involved in pathways regulating KCNQ channels**

| Candidate | Effect | Reference |
| --- | --- | --- |
| FGF14 | Positively regulates KCNQ channels | (Pablo & Pitt, 2017) |
| Kcne4 | Alters Vascular Reactivity through modulating KCNQ channels | (Abbott & Jepps, 2016) |
| PIP_2_ | Regulates KCNQ channel openings | (Zaydman et al., 2013 |
| cAMP/PKA | Enhance KCNQ currents | (Mani et al., 2016) |
| SGK-1 and Nedd4-2 | Modulates KCNQ channels by SGK-1 regulation of the activity of the ubiquitin ligase Nedd4-2 | (Schuetz, Kumar, Poronnik, & Adams, 2008) |
| SMIT1 or Slc5a3 | Regulates KCNQ channel ion selectivity | (Barrese, Stott, Baldwin, Mondejar-Parreno, & Greenwood, 2020; Manville, Neverisky, & Abbott, 2017) |

**Table S6. Significantly dysregulated genes in PVAT (RNA-Seq) and in white adipose tissue (WAT) (proteomics) in aging.**

| **Gene Symbol (mRNA, from current study)** | **Gene Symbol (proteomics from (Yu et al., 2020))** | **Gene description** | **Expression** | **Process** |
| --- | --- | --- | --- | --- |
| Abhd14b | Abhd14b | Abhydrolase domain containing 14b | ↓ | Lipid Metabolism |
| Abhd6 | Abhd6 | Abhydrolase domain containing 6 | ↓ |  |
| Acaca | Acaca | Acetyl-Coenzyme A carboxylase alpha | ↓ |  |
| Acacb | Acacb | Acetyl-Coenzyme A carboxylase beta | ↓ |  |
| Echs1 | Echs1 | Enoyl-CoA hydratase, mitochondrial | ↓ |  |
| Fasn | Fasn | Fatty acid synthase | ↓ |  |
| Gpd2 | Gpd2 | Pleckstrin homology domain-containing family O member 1 | ↓ |  |
| Acly | Acly | ATP citrate lyase | ↓ | Central Carbon |
| Gls | Gls | Glutaminase kidney isoform, mitochondrial | ↑ |  |
| Hk2 | Hk2 | Hexokinase-2 | ↓ |  |
| Mcee | Mcee | Methylmalonyl-CoA epimerase, mitochondrial | ↓ |  |
| Pdhb | Pdhb | Pyruvate dehydrogenase E1 component subunit beta, mitochondrial | ↓ |  |
| Pgk1 | Pgk1 | Phosphoglycerate kinase 1 | ↓ |  |
| Gpt2 | Gpt2 | Glutamic pyruvate transaminase | ↓ |  |
| Hk3 | Hk3 | Hexokinase-3 | ↑ |  |
| Cox5b |  | Cytochrome c oxidase subunit 5B, mitochondrial | ↓ | Electron Transport Chain |
| Cox6b1 | Cox6b1 | Cytochrome c oxidase subunit 6B1 | ↓ |  |
| Cox6c | Cox6c | Cytochrome c oxidase subunit 6C | ↓ |  |
| Ndufa3 | Ndufa3 | NADH dehydrogenase [ubiquinone] 1 alpha subcomplex subunit 3 | ↓ |  |
| Ndufa4 | Ndufa4 | NADH dehydrogenase [ubiquinone] 1 alpha subcomplex subunit 4 | ↓ |  |
| Ndufa5 | Ndufa5 | NADH dehydrogenase [ubiquinone] 1 alpha subcomplex subunit 5 | ↓ |  |
| Ndufa6 | Ndufa6 | NADH dehydrogenase [ubiquinone] 1 alpha subcomplex subunit 6 | ↓ |  |
| Ndufa7 | Ndufa7 | NADH dehydrogenase [ubiquinone] 1 alpha subcomplex subunit 7 | ↓ |  |
| Ndufa8 | Ndufa8 | NADH dehydrogenase [ubiquinone] 1 alpha subcomplex subunit 8 | ↓ |  |
| Ndufa10 | Ndufa10 | NADH dehydrogenase [ubiquinone] 1 alpha subcomplex subunit 10 | ↓ |  |
| Ndufa11 | Ndufa11 | NADH dehydrogenase [ubiquinone] 1 alpha subcomplex subunit 11 | ↓ |  |
| Ndufa12 | Ndufa12 | NADH dehydrogenase [ubiquinone] 1 alpha subcomplex subunit 12 | ↓ |  |
| Ndufb7 | Ndufb7 | NADH dehydrogenase [ubiquinone] 1 beta subcomplex subunit 7 | ↓ |  |
| Ndufb10 | Ndufb10 | NADH dehydrogenase [ubiquinone] 1 beta subcomplex subunit 10 | ↓ |  |
| Ndufb11 | Ndufb11 | NADH dehydrogenase [ubiquinone] 1 beta subcomplex subunit 11 | ↓ |  |
| Ndufs2 | Ndufs2 | NADH dehydrogenase [ubiquinone] iron-sulfur protein 2 | ↓ |  |
| Ndufs4 | Ndufs4 | NADH dehydrogenase [ubiquinone] iron-sulfur protein 4 | ↓ |  |
| Ndufs5 | Ndufs5 | NADH dehydrogenase [ubiquinone] iron-sulfur protein 5 | ↓ |  |
| Ndufs6 | Ndufs6 | NADH dehydrogenase [ubiquinone] iron-sulfur protein 6 | ↓ |  |
| Ndufs7 | Ndufs7 | NADH dehydrogenase [ubiquinone] iron-sulfur protein 7 | ↓ |  |
| Ndufv1 | Ndufv1 | NADH dehydrogenase [ubiquinone] flavoprotein 1 | ↓ |  |
| Ndufv2 | Ndufv2 | NADH dehydrogenase [ubiquinone] flavoprotein 2 | ↓ |  |
| Uqcrfs1 | Uqcrfs1 | Ubiquinol-cytochrome c reductase, Rieske iron-sulfur polypeptide 1 | ↓ |  |
| Casp1 | Casp1 | Caspase-1 | ↑ | Inflammation |
| Cd68 | Cd68 | Macrosialin | ↑ |  |
| Mrc1 | Mrc1 | Macrophage mannose receptor 1 | ↑ |  |
| Itgam | Itgam | Integrin alpha-M | ↑ |  |
| Stat2 | Stat2 | Signal transducer and activator of transcription 2 | ↑ |  |
| Rnasel | Rnasel | 2-5A-dependent ribonuclease | ↑ |  |

**References:**

Abbott, G. W., & Jepps, T. A. (2016). Kcne4 Deletion Sex-Dependently Alters Vascular Reactivity. *J Vasc Res, 53*(3-4), 138-148. doi:10.1159/000449060

Barrese, V., Stott, J. B., Baldwin, S. N., Mondejar-Parreno, G., & Greenwood, I. A. (2020). SMIT (Sodium-Myo-Inositol Transporter) 1 Regulates Arterial Contractility Through the Modulation of Vascular Kv7 Channels. *Arterioscler Thromb Vasc Biol, 40*(10), 2468-2480. doi:10.1161/ATVBAHA.120.315096

Mani, B. K., Robakowski, C., Brueggemann, L. I., Cribbs, L. L., Tripathi, A., Majetschak, M., & Byron, K. L. (2016). Kv7.5 Potassium Channel Subunits Are the Primary Targets for PKA-Dependent Enhancement of Vascular Smooth Muscle Kv7 Currents. *Mol Pharmacol, 89*(3), 323-334. doi:10.1124/mol.115.101758

Manville, R. W., Neverisky, D. L., & Abbott, G. W. (2017). SMIT1 Modifies KCNQ Channel Function and Pharmacology by Physical Interaction with the Pore. *Biophys J, 113*(3), 613-626. doi:10.1016/j.bpj.2017.06.055

Pablo, J. L., & Pitt, G. S. (2017). FGF14 is a regulator of KCNQ2/3 channels. *Proc Natl Acad Sci U S A, 114*(1), 154-159. doi:10.1073/pnas.1610158114

Schuetz, F., Kumar, S., Poronnik, P., & Adams, D. J. (2008). Regulation of the voltage-gated K(+) channels KCNQ2/3 and KCNQ3/5 by serum- and glucocorticoid-regulated kinase-1. *Am J Physiol Cell Physiol, 295*(1), C73-80. doi:10.1152/ajpcell.00146.2008

Yu, Q., Xiao, H., Jedrychowski, M. P., Schweppe, D. K., Navarrete-Perea, J., Knott, J., . . . Gygi, S. P. (2020). Sample multiplexing for targeted pathway proteomics in aging mice. *Proc Natl Acad Sci U S A, 117*(18), 9723-9732. doi:10.1073/pnas.1919410117
